# Telotrons: spliceosomal introns composed of telomeric repeats

**DOI:** 10.64898/2026.11.07.765104

**Authors:** Alexander M. Kramer, Russ Corbett-Detig

## Abstract

Spliceosomal intron numbers vary widely across eukaryotes, but the only established mechanism for large bursts of intron gain is the spread of transposable introner elements. We describe telotrons, spliceosomal introns composed predominantly of telomeric repeats. In a survey of 10,310 eukaryotic genome assemblies, we identified 7,008 introns with >85% telomeric repeat coverage and an additional 44,423 with >50% to ≤85% coverage, in 25 genomes spanning haptophytes, dinoflagellates, a cryptophyte, stramenopiles, the apicomplexan *Eimeria*, and the nematode *Mesorhabditis*. Most telotrons carry GT–AG or GC–AG splice boundaries, and RNA-seq supports their splicing. Their repeat orientations are consistent with telomerase extension of broken DNA ends, and intron absence at corresponding positions in orthologs supports widespread *de novo* gain. In *Eimeria* and *Mesorhabditis*, telotrons instead occupy positions where orthologs carry ordinary introns, suggesting modification of existing introns. Telotrons thus point to telomerase as an endogenous driver of structural innovation through intron creation and remodeling.

## Introduction

The origins of eukaryotic genome complexity remain a fundamental question of evolutionary biology, and spliceosomal intron number, in particular, varies by orders of magnitude across eukaryotes. Most lineages gain few introns, while others acquire hundreds to thousands along a single branch (Carmel et al. 2007; Csuros et al. 2011). The only established mechanism for gain on this scale is the proliferation of introners, transposable elements that encode or co-opt splice signals (Huff et al. 2016; Worden et al. 2009; van der Burgt et al. 2012; Gozashti et al. 2022, 2025). Other proposed mechanisms have not explained comparable large-scale bursts of new intron creation (Yenerall and Zhou 2012).

DNA double-strand break repair offers another route to intron formation. In *Daphnia*, direct repeats flanking recently gained introns implicate repair of staggered breaks as a mechanism of intron formation (W. Li et al. 2009, 2014). Telomerase could supply new sequence during this repair. This reverse transcriptase extends DNA 3′ ends with short repeats copied from its RNA template, and it can also add internal telomeric arrays (Nergadze et al. 2004, 2007; Kinzig et al. 2024). Splicing such an insertion from a coding transcript could create an intron.

Here, we describe telotrons, spliceosomal introns composed predominantly of telomeric repeats, in seven divergent eukaryotic lineages. Their structure points to telomerase extension of broken DNA ends as a source of new and remodeled introns.

## Results

We screened the annotated introns of 10,310 eukaryotic genome assemblies from NCBI RefSeq, GenBank and the Tara Oceans project (Karsenti et al. 2011; Sunagawa et al. 2020), using the published telomere motif of each lineage (Methods). We identified 7,008 telotrons with >85% telomeric repeat coverage in 25 genomes, each containing at least five qualifying introns (Fig. 1, supplementary fig. S2, Methods). Lowering the threshold to >50% coverage to capture interrupted or degraded arrays added 44,423 loci, for 51,431 telotrons in total. These genomes span haptophytes, dinoflagellates, a cryptophyte, a chrysophyte, another stramenopile, the apicomplexan *Eimeria*, and the nematode *Mesorhabditis*. Their scattered distribution among relatives lacking detected telotrons suggests independent expansions.

**Figure 1.**
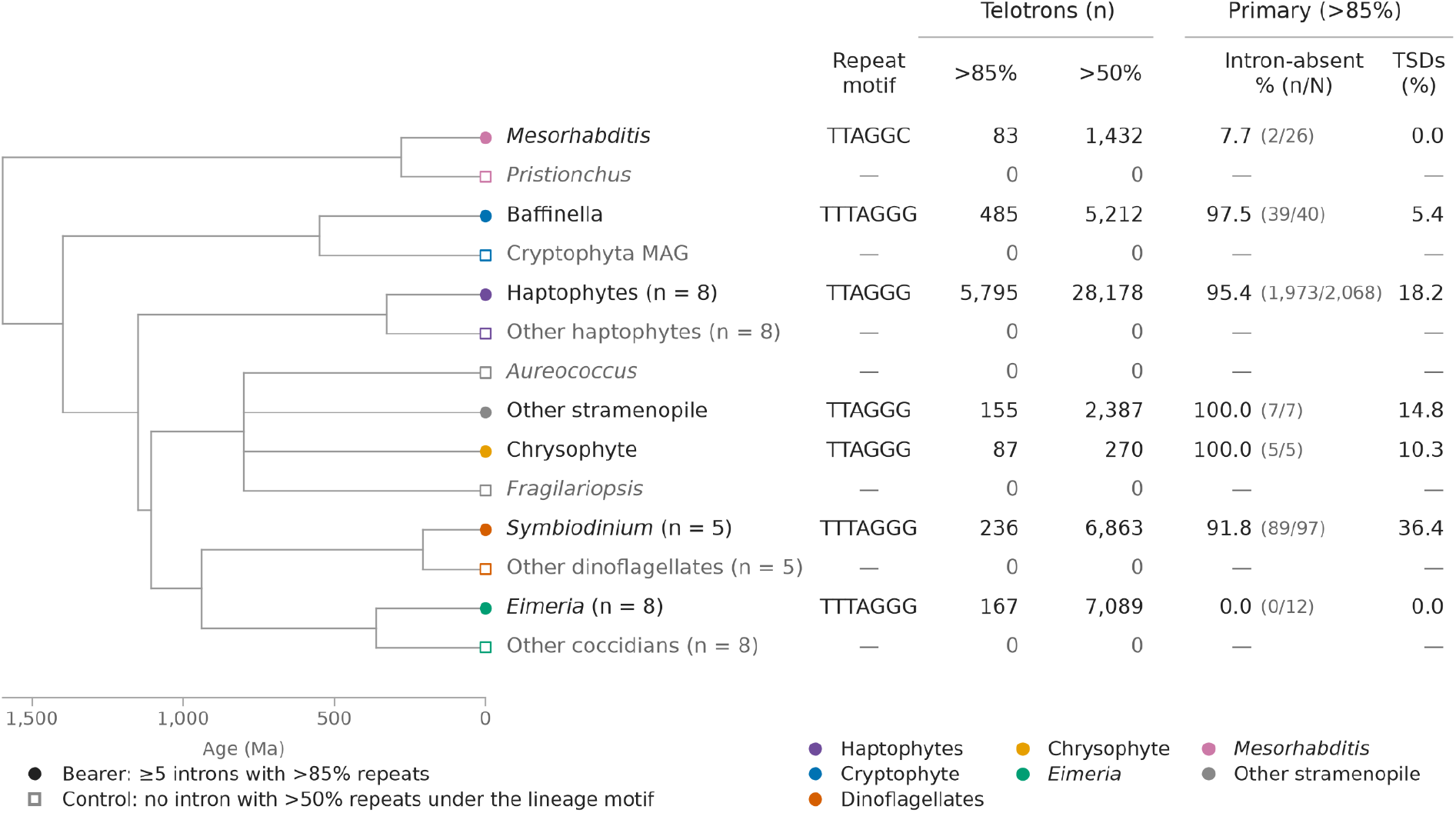
Distribution of telotrons across eukaryotes. Filled circles mark telotron bearer groups and open squares mark controls, with genome counts in parentheses for groups of more than one genome. Every genome was screened with the published telomere motif of its lineage (supplementary table S1), and controls have no intron above 50% repeat coverage under that motif. Colors indicate lineages. The >50% counts include the >85% set. Percentages of absent introns use primary loci whose closest qualifying comparison either lacks an intron or carries an ordinary intron supported by annotation at the same aligned position and phase (2,115/2,255 overall, 93.8%). TSD percentages use all primary loci and are pooled per lineage. Deep node ages follow the TimeTree backbone, and each lineage is drawn as an unresolved polytomy at a published crown age (Methods).

Telotrons carry recognizable splice signals, and RNA-seq supports their splicing. Among primary telotrons (>85% repeat coverage), 91.3% have GT–AG boundaries and 4.2% GC–AG, compared with 81.1% and 11.1% of additional telotrons (>50% to ≤85%). In the 11 genomes with RNA-seq data, junction reads supported splicing of 672 of 886 primary telotrons (75.8%) and 13,133 of 17,015 additional telotrons (77.2%), similar to ordinary introns in the same genomes under the same rule (68.0% of 22,000 sampled introns, supplementary table S3).

Short direct repeats at some telotron boundaries are consistent with insertion during DNA double-strand break repair (Fig. 2B). Gap filling during repair of staggered breaks can duplicate host sequence, producing target site duplications (TSDs), as proposed for other internal telomeric insertions (Nergadze et al. 2004; Santagostino et al. 2020). After excluding telomeric repeat sequence, we detected putative TSDs ≥6 bp at 1,200 of 7,008 primary telotrons (17.1%, Fig. 2B). None occurred in *Mesorhabditis* or *Eimeria*, frequencies in other lineages ranged from 5.4% to 36.4%, and lengths ranged from 6 to 28 bp (median 10 bp, supplementary table S2).

**Figure 2.**
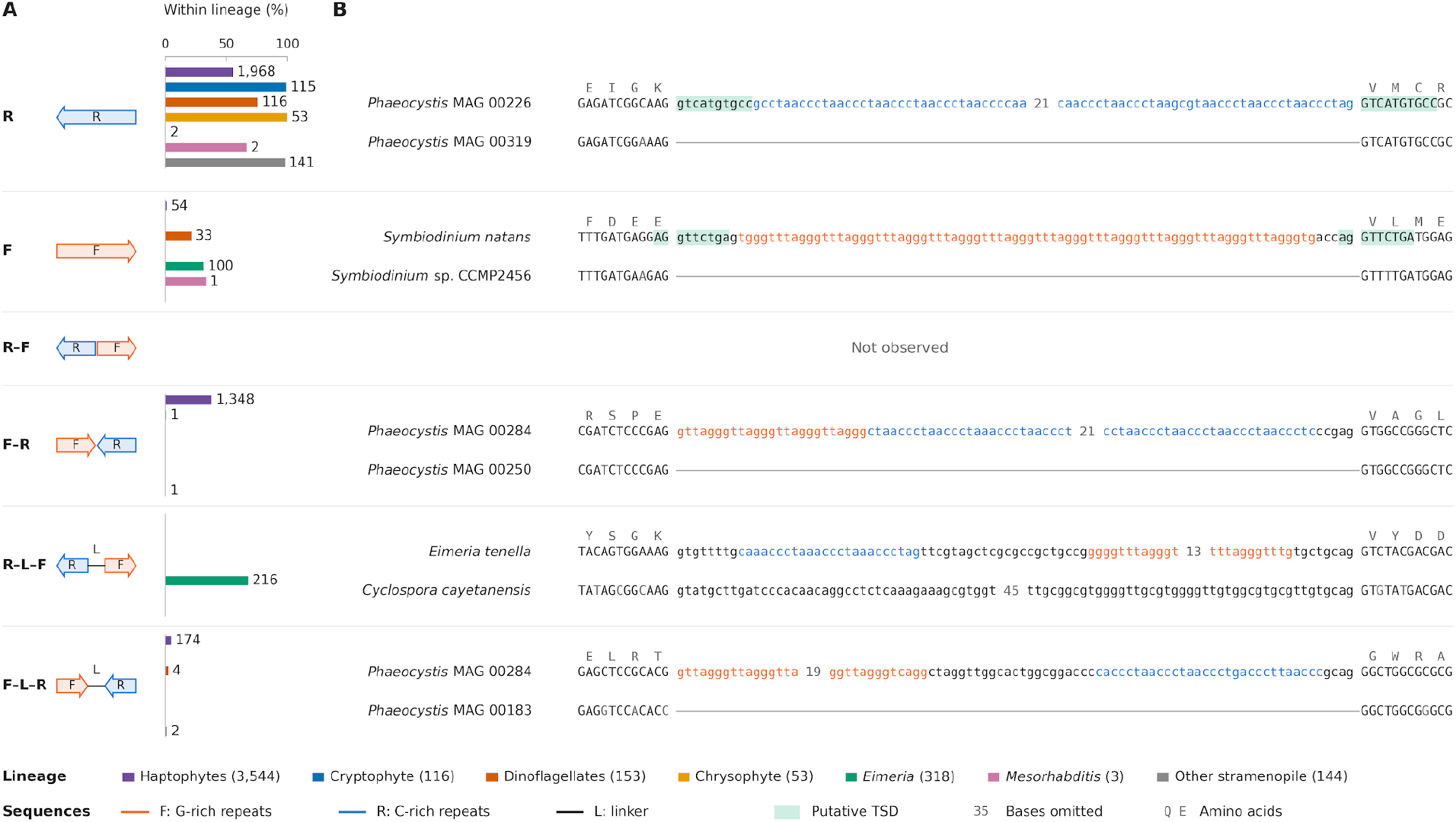
Telotron architecture and examples. (A) Architectures of 4,331 classified telotrons in 25 bearer genomes. Bars show percentages within each lineage among classified loci, labeled with counts. (B) Each example outside *Eimeria* shows the closest validated ortholog lacking the intron. *Eimeria* shows a comparison with an ordinary intron. Exons are uppercase and introns lowercase, and putative target site duplications (TSDs) are highlighted green. Gray letters show amino acids, numbers omitted bases and lines absent introns.

We classified 3,938 of 7,008 primary telotrons (56.2%) and 393 of 44,423 additional telotrons (0.9%) into five architectures (Fig. 2A). Along the transcript strand, F denotes G-rich arrays, R C-rich arrays and L non-telomeric linkers. Single C-rich arrays predominated (R, 2,397), followed by opposing arrays (F–R, 1,350) and single G-rich arrays (F, 188). No classified telotron had the two arrays directly adjacent in the reverse order (R–F). Telomerase extension of opposing 3′ ends predicts F before R, so this asymmetry is consistent with insertion by telomerase.

The remaining 396 classified telotrons had opposing arrays separated by a linker of at least 18 non-telomeric bases. Of these, 180 had the forward order (F–L–R) and 216 the reverse order (R–L–F). The reverse order therefore appeared only with a linker, and all 216 R–L–F telotrons were in *Eimeria*, with linkers of 19 to 109 bp (median 74 bp).

The absence of introns at corresponding positions in related species supports *de novo* telotron gain. The most closely related qualifying orthologs lacked the intron at 2,115 primary loci (93.8% of 2,255) and carried an ordinary intron at the same aligned position and phase at the other 140 (6.2%). In the five lineages other than *Eimeria* and *Mesorhabditis*, absence ranged from 91.8% to 100% (Fig. 1).

In contrast, ordinary non-telomeric introns occupied the same position and phase in all 12 informative *Eimeria* comparisons and in 24 of 26 *Mesorhabditis* comparisons, where the other two lacked an intron. These shared positions support modification of existing introns, although loss in relatives or independent gain at the same position could also contribute (Roy 2016; Klopfstein and Ronquist 2013; Torii et al. 2025; Collemare et al. 2015).

Bearer genomes also carried far more interstitial telomeric sequence (ITS) outside genes. Median ITS density was 665-fold higher in the 25 bearers than in 25 controls (177.8 versus 0.267 tracts per Mb, Fig. 3), and every bearer exceeded its paired control. Across seven selected lineage contrasts, bearers also exceeded their controls in non-coding ITS density (*W* = 0, *p* = 0.0156, two-sided Wilcoxon signed-rank test). This enrichment places telotrons within a broader accumulation of internal telomeric repeats, which telomerase can add during double-strand break repair (Nergadze et al. 2007; Kinzig et al. 2024), although it does not show that elevated ITS density alone produces telotrons.

**Figure 3.**
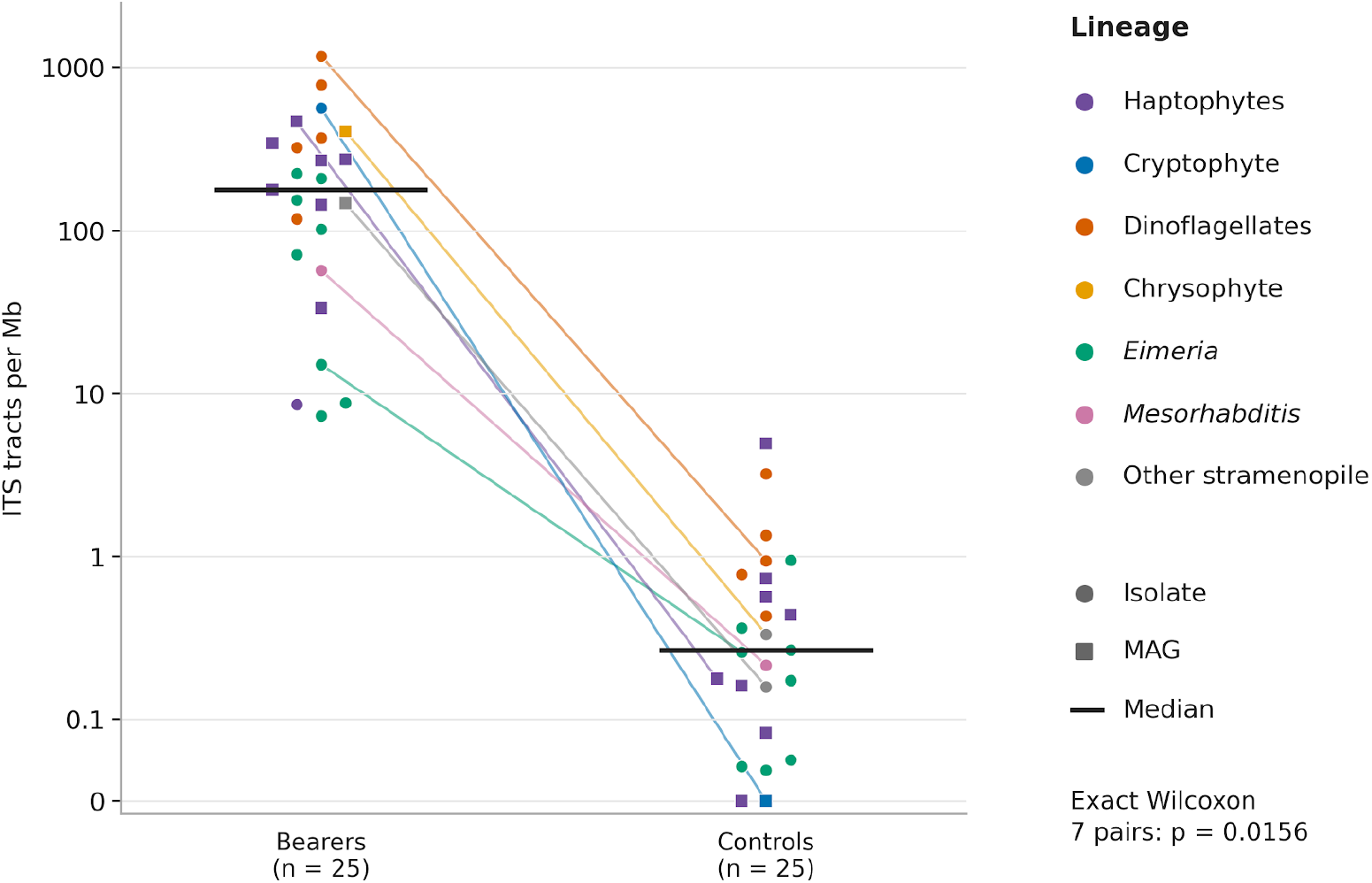
Interstitial telomeric sequence density in bearer genomes and controls. Each point represents one of 25 bearers or 25 distinct controls. Circles indicate isolates, squares MAGs and colors lineages. Black bars show medians of 177.8 and 0.267 tracts per Mb (665-fold difference). Tracts overlapping annotated or projected gene spans or terminal telomeric regions were removed. Density uses the remaining A/C/G/T sequence. Lines connect seven selected lineage pairs (exact two-sided Wilcoxon *W* = 0, *p* = 0.0156). The y-axis is logarithmic above 0.1 and linear below.

## Discussion

Both telomerase and the spliceosome date to early eukaryotic evolution. In one model, invading group II introns gave rise to both spliceosomal introns and linear chromosomes, and telomerase, likely descended from a retroelement reverse transcriptase, arose to maintain the new ends (Koonin 2006; Schrumpfová and Fajkus 2020). Intron generation by telomerase may therefore date to the earliest eukaryotes. Older telotrons would decay until they resembled ordinary introns, so we detect only those whose arrays remain telomeric.

About 0.2% of surveyed assemblies (25 of 10,310) carried telotrons, whereas we previously found recently active introners in 2.3% of 8,716 genomes (Gozashti et al. 2025). The surveys differ in sampling and detection methods, but recognizable telotrons appear less widespread than introners. Bearer genomes share a striking enrichment of interstitial telomeric sequence (Fig. 3), which suggests elevated rates of telomerase activity at breaks in these lineages, though what drives this remains unknown.

Our results support intron formation by telomerase extension of broken DNA ends, in line with experimental evidence that telomerase inserts telomeric repeats during double-strand break repair (Kinzig et al. 2024). F–R and F–L–R architectures place G-rich and C-rich arrays in the order expected from extension of opposing chromosomal 3′ ends, as proposed for internal telomeric arrays (Sola et al. 2021). F–L–R linkers would then be sequences retained or added during repair. The reversed R–L–F arrangement instead suggests that a free DNA fragment was extended and then inserted, with a linker of retained intron sequence or introduced DNA. Extrachromosomal fragments carrying fused telomeric arrays occur in human cells that use alternative lengthening of telomeres (Muyas et al. 2024), and capturing such fragments could also generate telotrons, but would need to explain why reversed arrays occur only with a linker. Experiments in a culturable telotron bearer could distinguish synthesis at breaks from capture of existing DNA.

Telotrons occur in at least three eukaryotic supergroups and reach thousands of copies in single genomes, revealing a source of genome-wide intron gain beyond introner proliferation. Telomerase maintains chromosome ends, and our findings suggest that it also creates and remodels spliceosomal introns.

## Methods

### Genomes and telotron identification

We obtained 10,310 assemblies and annotations from NCBI RefSeq, GenBank and Tara Oceans SMAGs-v1 (Delmont et al. 2022) and extracted introns between annotated exons or, when unavailable, CDS blocks. After an initial survey, we reanalyzed 45 candidate bearers and annotated comparison genomes (supplementary table S1). We retained 25 bearers with at least five primary loci and excluded eight with fewer. In Fig. 1, ages of nodes between lineages are TimeTree estimates (Kumar et al. 2022). Lineages are drawn as polytomies at a published crown age, with Rhabditina at 280 Ma (Dieterich et al. 2008), Prymnesiophyceae 329 Ma (Liu et al. 2010), Eimeriorina 361 Ma (Hong et al. 2026), Cryptophyta 550 Ma (Pietluch et al. 2024), Ochrophyta 800 Ma (Choi et al. 2024) and Suessiales 207 Ma, from Late Triassic Suessiaceae cysts (Montresor et al. 1999).

We scanned each lineage’s published telomere motif (supplementary table S1) in all rotations and both orientations. Coverage was the fraction of intron bases in exact units or two-unit windows with ≤1 substitution. Introns covered >85% by any other repeat were comparatively rare in bearers and controls (supplementary fig. S3).

We required introns ≥30 bp with no splice signal filter. We excluded candidates with ≥50% repeat coverage in either adjacent 12-bp flank or >20% across flanks of up to 200 bp per side. Primary telotrons had >85% coverage and additional telotrons >50% to ≤85%.

We grouped loci whose flanks were both aligned by BLASTn (Camacho et al. 2009) over ≥100 bp at ≥95% identity and kept the longest primary telotron per group, or the longest intron if none qualified.

### Repeat architecture

We classified arrays on the transcript strand as G-rich (F) or C-rich (R), each with at least three motif units, merging same-orientation arrays across gaps of ≤3 bp. Both outer endpoints had to fall within 8 bp of the intron ends, or of the inner edge of the intronic copy of a putative TSD. Degraded repeats extending in phase from an array for at least three units with up to two substitutions per unit were masked before measuring linkers.

Directly opposing arrays formed F–R or R–F configurations. Linker architectures required ≥18 consecutive repeat-free bases between arrays and >50% coverage, and architectures without a linker required >85%. Other loci were left unclassified (supplementary fig. S1).

### Target site duplications

We searched the intron and up to 60 bp of each adjoining exon, outward from the junctions, for exact non-telomeric direct repeats spanning both splice junctions. We required a combined match ≥6 bp with at least 4 bp beyond the two junction-adjacent bases and excluded single-base and low-complexity matches.

### RNA-seq support

We aligned public RNA-seq reads from 11 genomes (supplementary table S3) with minimap2 (H. Li 2018) in splice:sr mode and counted a locus as supported by at least one primary alignment with MAPQ ≥20 spanning the exact annotated junction with ≥10 contiguous aligned bases on each side. As a baseline we applied the same rule to a random sample of 2,000 ordinary introns per genome (≥30 bp, <30% telomeric coverage).

### Ortholog comparisons

We mapped host proteins to comparator genomes with miniprot (H. Li 2023). For each bearer we chose up to 40 markers, the longest annotated proteins of 200 to 1,500 aa with no internal stop codon. We mapped them to every candidate genome, ranked candidates by mean marker identity and recovery, and then added every candidate in which miniprot recovered the host protein segment spanning the telotron. We dropped comparisons in which a second genomic hit covered ≥90% of the best hit’s span.

We required reciprocal best hits, ≥40% whole-protein identity and ≥12 matches in each 20-residue junction flank, and assessed positional correspondence with MAFFT protein alignments (Katoh and Standley 2013). An ordinary intron had to match the aligned coding position and phase, agree with an annotated coding model, and have <30% telomeric coverage with no tract of ≥4 units.

For each locus we used the qualifying comparison with the highest identity, whether the intron was absent or ordinary, left ties between states unresolved, and calculated percentages across supported comparisons.

### ITS density and controls

We detected telomeric tracts of ≥4 units with ≤1 substitution per adjacent pair and removed tracts overlapping annotated genes, terminal telomeric regions or gene spans projected by miniprot. Density was the remaining tract count per megabase of unmasked A/C/G/T sequence. Contig ends were masked inward while the lineage motif covered ≥15% of each 1-kb window. Projected genes were miniprot alignments of same-lineage proteins from other genomes (≥50% of the protein at ≥40% identity over ≥100 residues).

We selected 25 controls within lineages by annotation quality and marker-tree distance, each screened with its bearers’ lineage motif and with no intron above 50% repeat coverage (supplementary table S1).

We compared ITS densities across seven selected bearer–control pairs using an exact two-sided Wilcoxon signed-rank test.

## Data and code availability

Code for every analysis and figure is available at github.com/amkram/telotrons. The catalogue of all telotrons with evidence for each locus is in supplementary table S2. Genome assemblies, annotations and RNA-seq runs are public, with accessions in supplementary tables S1 and S3.

## Funding Acknowledgement

This work was funded by R35GM128932 to RC-D.

## AI Acknowledgement

AI tools were used to develop code and for proofreading text. All AI-generated elements have been reviewed by the authors.

## Acknowledgements

The authors thank Manny Ares and Carol Greider for helpful feedback on initial results in this study.

## SUPPLEMENTARY INFORMATION

### Supplementary Tables

**Supplementary Table S1.**
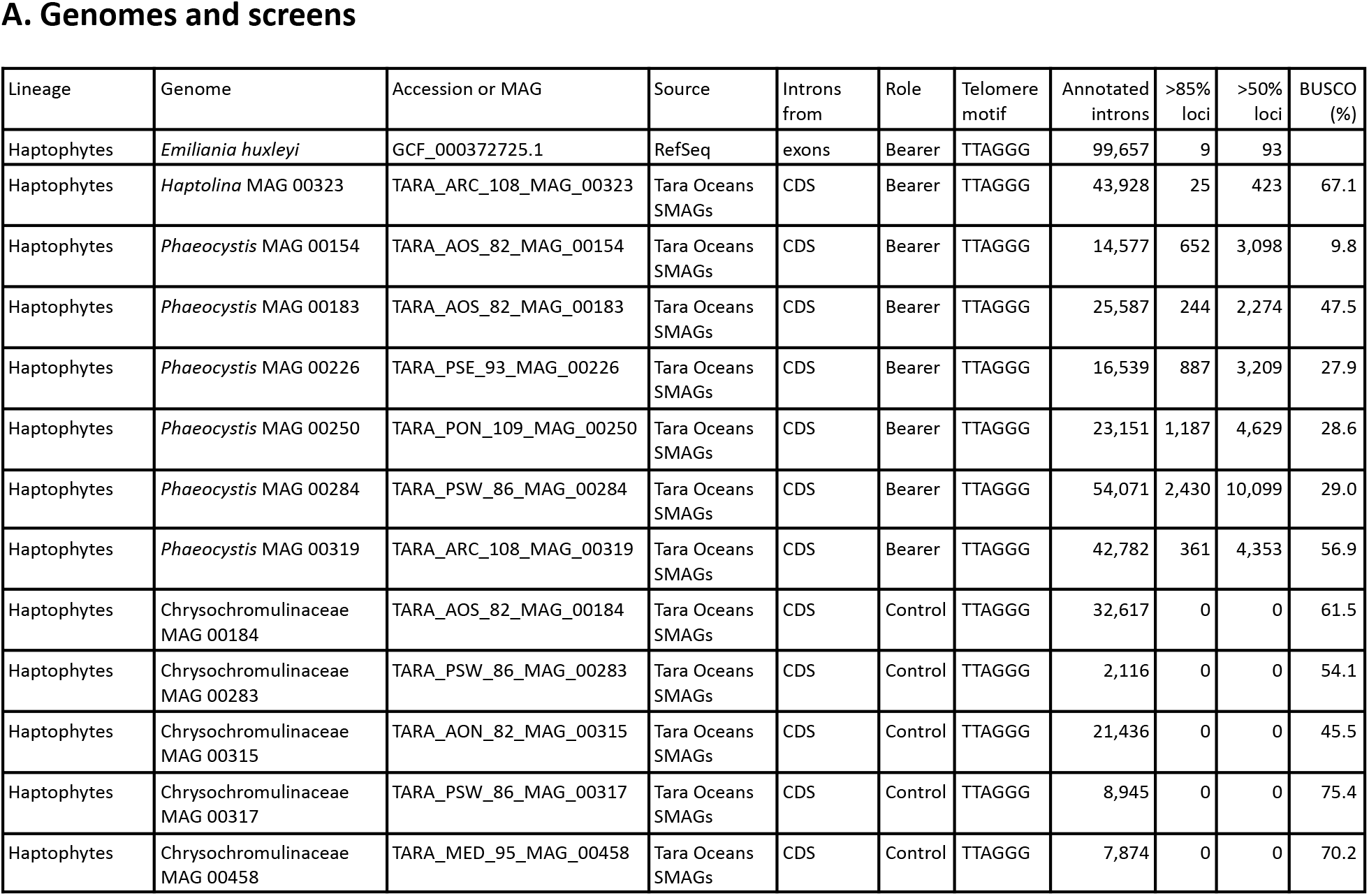
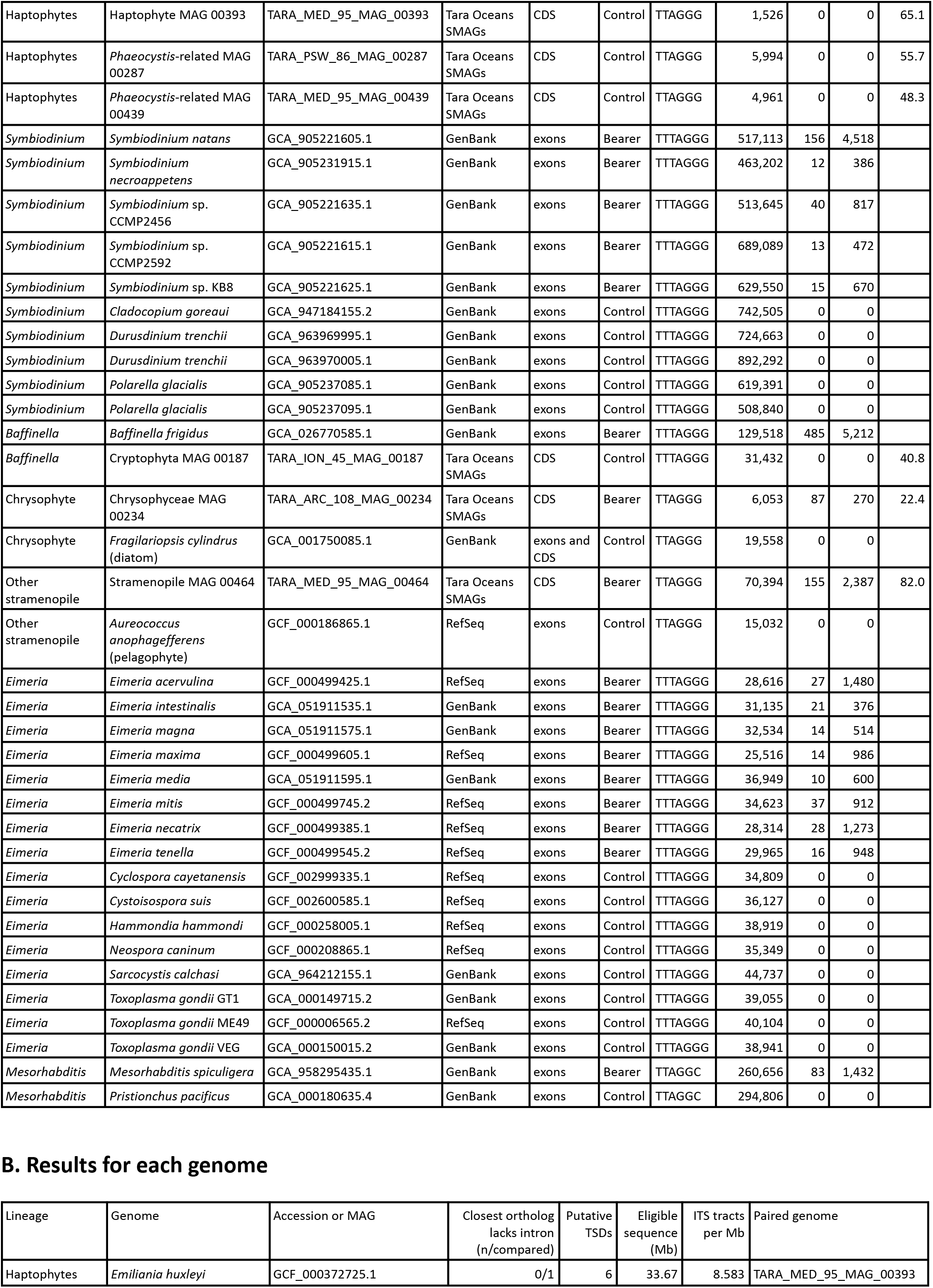
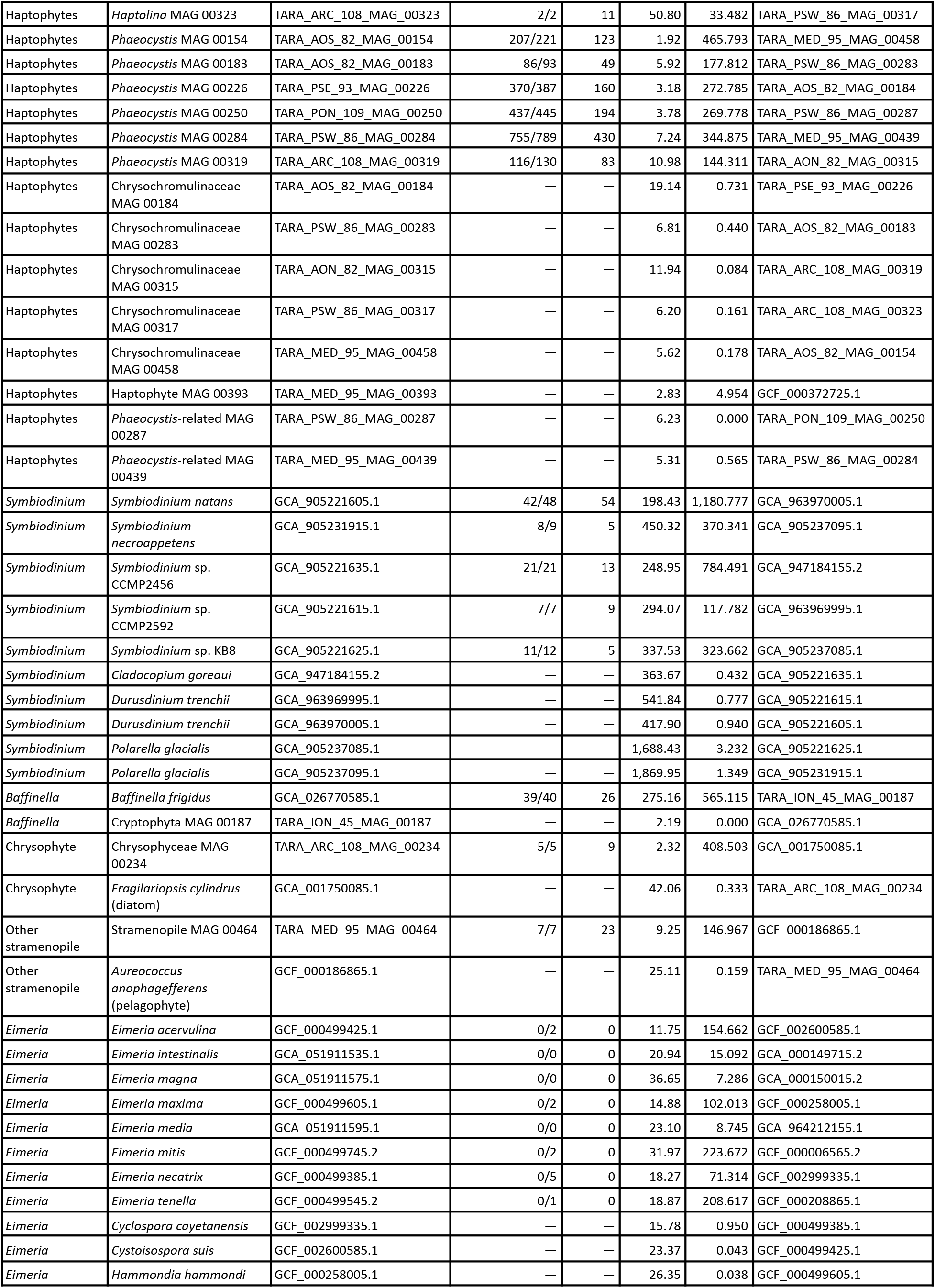
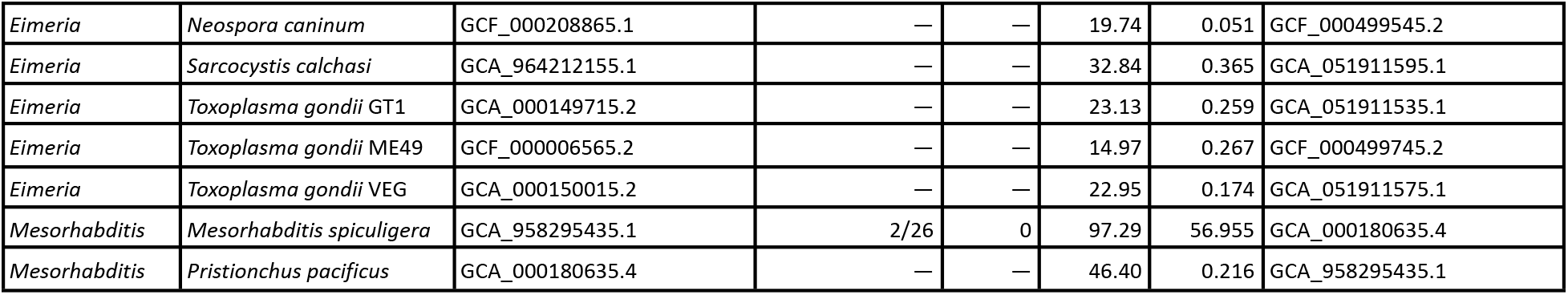
Genome identity, screen status and results for each genome. Part A identifies each genome. BUSCO completeness of MAGs is from the SMAG release (Delmont et al. 2022). Each genome was screened with the published telomere motif of its lineage in all cyclic rotations and both orientations. Part B gives results for each genome. The ortholog column counts primary loci whose closest qualifying ortholog lacks the intron, over loci with a qualifying comparison (intron absent, or an ordinary intron supported by annotation at the same position and phase). The TSD column counts putative target site duplications at primary loci. The last three columns give the eligible sequence length, the density of interstitial telomeric sequence (ITS) tracts outside genes, and the paired genome used in Fig. 3.

| Lineage | Genome | Accession or MAG | Source | Introns from | Role | Telomere motif | Annotated introns | >85% loci | >50% loci | BUSCO (%) |
|---|---|---|---|---|---|---|---|---|---|---|
| Haptophytes | *Emiliania huxleyi* | GCF_000372725.1 | RefSeq | exons | Bearer | TTAGGG | 99,657 | 9 | 93 |  |
| Haptophytes | *Haptolina* MAG 00323 | TARA_ARC_108_MAG_00323 | Tara Oceans SMAGs | CDS | Bearer | TTAGGG | 43,928 | 25 | 423 | 67.1 |
| Haptophytes | *Phaeocystis* MAG 00154 | TARA_AOS_82_MAG_00154 | Tara Oceans SMAGs | CDS | Bearer | TTAGGG | 14,577 | 652 | 3,098 | 9.8 |
| Haptophytes | *Phaeocystis* MAG 00183 | TARA_AOS_82_MAG_00183 | Tara Oceans SMAGs | CDS | Bearer | TTAGGG | 25,587 | 244 | 2,274 | 47.5 |
| Haptophytes | *Phaeocystis* MAG 00226 | TARA_PSE_93_MAG_00226 | Tara Oceans SMAGs | CDS | Bearer | TTAGGG | 16,539 | 887 | 3,209 | 27.9 |
| Haptophytes | *Phaeocystis* MAG 00250 | TARA_PON_109_MAG_00250 | Tara Oceans SMAGs | CDS | Bearer | TTAGGG | 23,151 | 1,187 | 4,629 | 28.6 |
| Haptophytes | *Phaeocystis* MAG 00284 | TARA_PSW_86_MAG_00284 | Tara Oceans SMAGs | CDS | Bearer | TTAGGG | 54,071 | 2,430 | 10,099 | 29.0 |
| Haptophytes | *Phaeocystis* MAG 00319 | TARA_ARC_108_MAG_00319 | Tara Oceans SMAGs | CDS | Bearer | TTAGGG | 42,782 | 361 | 4,353 | 56.9 |
| Haptophytes | Chrysochromulinaceae MAG 00184 | TARA_AOS_82_MAG_00184 | Tara Oceans SMAGs | CDS | Control | TTAGGG | 32,617 | 0 | 0 | 61.5 |
| Haptophytes | Chrysochromulinaceae MAG 00283 | TARA_PSW_86_MAG_00283 | Tara Oceans SMAGs | CDS | Control | TTAGGG | 2,116 | 0 | 0 | 54.1 |
| Haptophytes | Chrysochromulinaceae MAG 00315 | TARA_AON_82_MAG_00315 | Tara Oceans SMAGs | CDS | Control | TTAGGG | 21,436 | 0 | 0 | 45.5 |
| Haptophytes | Chrysochromulinaceae MAG 00317 | TARA_PSW_86_MAG_00317 | Tara Oceans SMAGs | CDS | Control | TTAGGG | 8,945 | 0 | 0 | 75.4 |
| Haptophytes | Chrysochromulinaceae MAG 00458 | TARA_MED_95_MAG_00458 | Tara Oceans SMAGs | CDS | Control | TTAGGG | 7,874 | 0 | 0 | 70.2 |
| Haptophytes | Haptophyte MAG 00393 | TARA_MED_95_MAG_00393 | Tara Oceans SMAGs | CDS | Control | TTAGGG | 1,526 | 0 | 0 | 65.1 |
| Haptophytes | *Phaeocystis*-related MAG 00287 | TARA_PSW_86_MAG_00287 | Tara Oceans SMAGs | CDS | Control | TTAGGG | 5,994 | 0 | 0 | 55.7 |
| Haptophytes | *Phaeocystis*-related MAG 00439 | TARA_MED_95_MAG_00439 | Tara Oceans SMAGs | CDS | Control | TTAGGG | 4,961 | 0 | 0 | 48.3 |
| *Symbiodinium* | *Symbiodinium natans* | GCA_905221605.1 | GenBank | exons | Bearer | TTTAGGG | 517,113 | 156 | 4,518 |  |
| *Symbiodinium* | *Symbiodinium necroappetens* | GCA_905231915.1 | GenBank | exons | Bearer | TTTAGGG | 463,202 | 12 | 386 |  |
| *Symbiodinium* | *Symbiodinium* sp. CCMP2456 | GCA_905221635.1 | GenBank | exons | Bearer | TTTAGGG | 513,645 | 40 | 817 |  |
| *Symbiodinium* | *Symbiodinium* sp. CCMP2592 | GCA_905221615.1 | GenBank | exons | Bearer | TTTAGGG | 689,089 | 13 | 472 |  |
| *Symbiodinium* | *Symbiodinium* sp. KB8 | GCA_905221625.1 | GenBank | exons | Bearer | TTTAGGG | 629,550 | 15 | 670 |  |
| *Symbiodinium* | *Cladocopium goreaui* | GCA_947184155.2 | GenBank | exons | Control | TTTAGGG | 742,505 | 0 | 0 |  |
| *Symbiodinium* | *Durusdinium trenchii* | GCA_963969995.1 | GenBank | exons | Control | TTTAGGG | 724,663 | 0 | 0 |  |
| *Symbiodinium* | *Durusdinium trenchii* | GCA_963970005.1 | GenBank | exons | Control | TTTAGGG | 892,292 | 0 | 0 |  |
| *Symbiodinium* | *Polarella glacialis* | GCA_905237085.1 | GenBank | exons | Control | TTTAGGG | 619,391 | 0 | 0 |  |
| *Symbiodinium* | *Polarella glacialis* | GCA_905237095.1 | GenBank | exons | Control | TTTAGGG | 508,840 | 0 | 0 |  |
| *Baffinella* | *Baffinella frigidus* | GCA_026770585.1 | GenBank | exons | Bearer | TTTAGGG | 129,518 | 485 | 5,212 |  |
| *Baffinella* | Cryptophyta MAG 00187 | TARA_ION_45_MAG_00187 | Tara Oceans SMAGs | CDS | Control | TTTAGGG | 31,432 | 0 | 0 | 40.8 |
| Chrysophyte | Chrysophyceae MAG 00234 | TARA_ARC_108_MAG_00234 | Tara Oceans SMAGs | CDS | Bearer | TTAGGG | 6,053 | 87 | 270 | 22.4 |
| Chrysophyte | *Fragilariopsis cylindrus* (diatom) | GCA_001750085.1 | GenBank | exons and CDS | Control | TTAGGG | 19,558 | 0 | 0 |  |
| Other stramenopile | Stramenopile MAG 00464 | TARA_MED_95_MAG_00464 | Tara Oceans SMAGs | CDS | Bearer | TTAGGG | 70,394 | 155 | 2,387 | 82.0 |
| Other stramenopile | *Aureococcus anophagefferens* (pelagophyte) | GCF_000186865.1 | RefSeq | exons | Control | TTAGGG | 15,032 | 0 | 0 |  |
| *Eimeria* | *Eimeria acervulina* | GCF_000499425.1 | RefSeq | exons | Bearer | TTTAGGG | 28,616 | 27 | 1,480 |  |
| *Eimeria* | *Eimeria intestinalis* | GCA_051911535.1 | GenBank | exons | Bearer | TTTAGGG | 31,135 | 21 | 376 |  |
| *Eimeria* | *Eimeria magna* | GCA_051911575.1 | GenBank | exons | Bearer | TTTAGGG | 32,534 | 14 | 514 |  |
| *Eimeria* | *Eimeria maxima* | GCF_000499605.1 | RefSeq | exons | Bearer | TTTAGGG | 25,516 | 14 | 986 |  |
| *Eimeria* | *Eimeria media* | GCA_051911595.1 | GenBank | exons | Bearer | TTTAGGG | 36,949 | 10 | 600 |  |
| *Eimeria* | *Eimeria mitis* | GCF_000499745.2 | RefSeq | exons | Bearer | TTTAGGG | 34,623 | 37 | 912 |  |
| *Eimeria* | *Eimeria necatrix* | GCF_000499385.1 | RefSeq | exons | Bearer | TTTAGGG | 28,314 | 28 | 1,273 |  |
| *Eimeria* | *Eimeria tenella* | GCF_000499545.2 | RefSeq | exons | Bearer | TTTAGGG | 29,965 | 16 | 948 |  |
| *Eimeria* | *Cyclospora cayetanensis* | GCF_002999335.1 | RefSeq | exons | Control | TTTAGGG | 34,809 | 0 | 0 |  |
| *Eimeria* | *Cystoisospora suis* | GCF_002600585.1 | RefSeq | exons | Control | TTTAGGG | 36,127 | 0 | 0 |  |
| *Eimeria* | *Hammondia hammondi* | GCF_000258005.1 | RefSeq | exons | Control | TTTAGGG | 38,919 | 0 | 0 |  |
| *Eimeria* | *Neospora caninum* | GCF_000208865.1 | RefSeq | exons | Control | TTTAGGG | 35,349 | 0 | 0 |  |
| *Eimeria* | *Sarcocystis calchasi* | GCA_964212155.1 | GenBank | exons | Control | TTTAGGG | 44,737 | 0 | 0 |  |
| *Eimeria* | *Toxoplasma gondii* GT1 | GCA_000149715.2 | GenBank | exons | Control | TTTAGGG | 39,055 | 0 | 0 |  |
| *Eimeria* | *Toxoplasma gondii* ME49 | GCF_000006565.2 | RefSeq | exons | Control | TTTAGGG | 40,104 | 0 | 0 |  |
| *Eimeria* | *Toxoplasma gondii* VEG | GCA_000150015.2 | GenBank | exons | Control | TTTAGGG | 38,941 | 0 | 0 |  |
| *Mesorhabditis* | *Mesorhabditis spiculigera* | GCA_958295435.1 | GenBank | exons | Bearer | TTAGGC | 260,656 | 83 | 1,432 |  |
| *Mesorhabditis* | *Pristionchus pacificus* | GCA_000180635.4 | GenBank | exons | Control | TTAGGC | 294,806 | 0 | 0 |  |
| Lineage | Genome | Accession or MAG | Closest ortholog lacks intron (n/compared) | Putative TSDs | Eligible sequence (Mb) | ITS tracts per Mb | Paired genome |  |  |  |
| Haptophytes | *Emiliania huxleyi* | GCF_000372725.1 | 0/1 | 6 | 33.67 | 8.583 | TARA_MED_95_MAG_00393 |  |  |  |
| Haptophytes | *Haptolina* MAG 00323 | TARA_ARC_108_MAG_00323 | 2/2 | 11 | 50.80 | 33.482 | TARA_PSW_86_MAG_00317 |  |  |  |
| Haptophytes | *Phaeocystis* MAG 00154 | TARA_AOS_82_MAG_00154 | 207/221 | 123 | 1.92 | 465.793 | TARA_MED_95_MAG_00458 |  |  |  |
| Haptophytes | *Phaeocystis* MAG 00183 | TARA_AOS_82_MAG_00183 | 86/93 | 49 | 5.92 | 177.812 | TARA_PSW_86_MAG_00283 |  |  |  |
| Haptophytes | *Phaeocystis* MAG 00226 | TARA_PSE_93_MAG_00226 | 370/387 | 160 | 3.18 | 272.785 | TARA_AOS_82_MAG_00184 |  |  |  |
| Haptophytes | *Phaeocystis* MAG 00250 | TARA_PON_109_MAG_00250 | 437/445 | 194 | 3.78 | 269.778 | TARA_PSW_86_MAG_00287 |  |  |  |
| Haptophytes | *Phaeocystis* MAG 00284 | TARA_PSW_86_MAG_00284 | 755/789 | 430 | 7.24 | 344.875 | TARA_MED_95_MAG_00439 |  |  |  |
| Haptophytes | *Phaeocystis* MAG 00319 | TARA_ARC_108_MAG_00319 | 116/130 | 83 | 10.98 | 144.311 | TARA_AON_82_MAG_00315 |  |  |  |
| Haptophytes | Chrysochromulinaceae MAG 00184 | TARA_AOS_82_MAG_00184 | — | — | 19.14 | 0.731 | TARA_PSE_93_MAG_00226 |  |  |  |
| Haptophytes | Chrysochromulinaceae MAG 00283 | TARA_PSW_86_MAG_00283 | — | — | 6.81 | 0.440 | TARA_AOS_82_MAG_00183 |  |  |  |
| Haptophytes | Chrysochromulinaceae MAG 00315 | TARA_AON_82_MAG_00315 | — | — | 11.94 | 0.084 | TARA_ARC_108_MAG_00319 |  |  |  |
| Haptophytes | Chrysochromulinaceae MAG 00317 | TARA_PSW_86_MAG_00317 | — | — | 6.20 | 0.161 | TARA_ARC_108_MAG_00323 |  |  |  |
| Haptophytes | Chrysochromulinaceae MAG 00458 | TARA_MED_95_MAG_00458 | — | — | 5.62 | 0.178 | TARA_AOS_82_MAG_00154 |  |  |  |
| Haptophytes | Haptophyte MAG 00393 | TARA_MED_95_MAG_00393 | — | — | 2.83 | 4.954 | GCF_000372725.1 |  |  |  |
| Haptophytes | *Phaeocystis*-related MAG 00287 | TARA_PSW_86_MAG_00287 | — | — | 6.23 | 0.000 | TARA_PON_109_MAG_00250 |  |  |  |
| Haptophytes | *Phaeocystis*-related MAG 00439 | TARA_MED_95_MAG_00439 | — | — | 5.31 | 0.565 | TARA_PSW_86_MAG_00284 |  |  |  |
| *Symbiodinium* | *Symbiodinium natans* | GCA_905221605.1 | 42/48 | 54 | 198.43 | 1,180.777 | GCA_963970005.1 |  |  |  |
| *Symbiodinium* | *Symbiodinium necroappetens* | GCA_905231915.1 | 8/9 | 5 | 450.32 | 370.341 | GCA_905237095.1 |  |  |  |
| *Symbiodinium* | *Symbiodinium* sp. CCMP2456 | GCA_905221635.1 | 21/21 | 13 | 248.95 | 784.491 | GCA_947184155.2 |  |  |  |
| *Symbiodinium* | *Symbiodinium* sp. CCMP2592 | GCA_905221615.1 | 7/7 | 9 | 294.07 | 117.782 | GCA_963969995.1 |  |  |  |
| *Symbiodinium* | *Symbiodinium* sp. KB8 | GCA_905221625.1 | 11/12 | 5 | 337.53 | 323.662 | GCA_905237085.1 |  |  |  |
| *Symbiodinium* | *Cladocopium goreaui* | GCA_947184155.2 | — | — | 363.67 | 0.432 | GCA_905221635.1 |  |  |  |
| *Symbiodinium* | *Durusdinium trenchii* | GCA_963969995.1 | — | — | 541.84 | 0.777 | GCA_905221615.1 |  |  |  |
| *Symbiodinium* | *Durusdinium trenchii* | GCA_963970005.1 | — | — | 417.90 | 0.940 | GCA_905221605.1 |  |  |  |
| *Symbiodinium* | *Polarella glacialis* | GCA_905237085.1 | — | — | 1,688.43 | 3.232 | GCA_905221625.1 |  |  |  |
| *Symbiodinium* | *Polarella glacialis* | GCA_905237095.1 | — | — | 1,869.95 | 1.349 | GCA_905231915.1 |  |  |  |
| *Baffinella* | *Baffinella frigidus* | GCA_026770585.1 | 39/40 | 26 | 275.16 | 565.115 | TARA_ION_45_MAG_00187 |  |  |  |
| *Baffinella* | Cryptophyta MAG 00187 | TARA_ION_45_MAG_00187 | — | — | 2.19 | 0.000 | GCA_026770585.1 |  |  |  |
| Chrysophyte | Chrysophyceae MAG 00234 | TARA_ARC_108_MAG_00234 | 5/5 | 9 | 2.32 | 408.503 | GCA_001750085.1 |  |  |  |
| Chrysophyte | *Fragilariopsis cylindrus* (diatom) | GCA_001750085.1 | — | — | 42.06 | 0.333 | TARA_ARC_108_MAG_00234 |  |  |  |
| Other stramenopile | Stramenopile MAG 00464 | TARA_MED_95_MAG_00464 | 7/7 | 23 | 9.25 | 146.967 | GCF_000186865.1 |  |  |  |
| Other stramenopile | *Aureococcus anophagefferens* (pelagophyte) | GCF_000186865.1 | — | — | 25.11 | 0.159 | TARA_MED_95_MAG_00464 |  |  |  |
| *Eimeria* | *Eimeria acervulina* | GCF_000499425.1 | 0/2 | 0 | 11.75 | 154.662 | GCF_002600585.1 |  |  |  |
| *Eimeria* | *Eimeria intestinalis* | GCA_051911535.1 | 0/0 | 0 | 20.94 | 15.092 | GCA_000149715.2 |  |  |  |
| *Eimeria* | *Eimeria magna* | GCA_051911575.1 | 0/0 | 0 | 36.65 | 7.286 | GCA_000150015.2 |  |  |  |
| *Eimeria* | *Eimeria maxima* | GCF_000499605.1 | 0/2 | 0 | 14.88 | 102.013 | GCF_000258005.1 |  |  |  |
| *Eimeria* | *Eimeria media* | GCA_051911595.1 | 0/0 | 0 | 23.10 | 8.745 | GCA_964212155.1 |  |  |  |
| *Eimeria* | *Eimeria mitis* | GCF_000499745.2 | 0/2 | 0 | 31.97 | 223.672 | GCF_000006565.2 |  |  |  |
| *Eimeria* | *Eimeria necatrix* | GCF_000499385.1 | 0/5 | 0 | 18.27 | 71.314 | GCF_002999335.1 |  |  |  |
| *Eimeria* | *Eimeria tenella* | GCF_000499545.2 | 0/1 | 0 | 18.87 | 208.617 | GCF_000208865.1 |  |  |  |
| *Eimeria* | *Cyclospora cayetanensis* | GCF_002999335.1 | — | — | 15.78 | 0.950 | GCF_000499385.1 |  |  |  |
| *Eimeria* | *Cystoisospora suis* | GCF_002600585.1 | — | — | 23.37 | 0.043 | GCF_000499425.1 |  |  |  |
| *Eimeria* | *Hammondia hammondi* | GCF_000258005.1 | — | — | 26.35 | 0.038 | GCF_000499605.1 |  |  |  |
| *Eimeria* | *Neospora caninum* | GCF_000208865.1 | — | — | 19.74 | 0.051 | GCF_000499545.2 |  |  |  |
| *Eimeria* | *Sarcocystis calchasi* | GCA_964212155.1 | — | — | 32.84 | 0.365 | GCA_051911595.1 |  |  |  |
| *Eimeria* | *Toxoplasma gondii* GT1 | GCA_000149715.2 | — | — | 23.13 | 0.259 | GCA_051911535.1 |  |  |  |
| *Eimeria* | *Toxoplasma gondii* ME49 | GCF_000006565.2 | — | — | 14.97 | 0.267 | GCF_000499745.2 |  |  |  |
| *Eimeria* | *Toxoplasma gondii* VEG | GCA_000150015.2 | — | — | 22.95 | 0.174 | GCA_051911575.1 |  |  |  |
| *Mesorhabditis* | *Mesorhabditis spiculigera* | GCA_958295435.1 | 2/26 | 0 | 97.29 | 56.955 | GCA_000180635.4 |  |  |  |
| *Mesorhabditis* | *Pristionchus pacificus* | GCA_000180635.4 | — | — | 46.40 | 0.216 | GCA_958295435.1 |  |  |  |

**Supplementary Table S2. Telotron catalogue with evidence for each locus**. The full catalogue of 51,431 telotrons is a separate spreadsheet (SUPPLEMENTARY_TABLES.xlsx). Each row gives coordinates, strand, intron length, splice boundaries, repeat coverage, architecture or the first rule an unclassified locus failed, linker coordinates, the putative TSD, RNA junction reads, the state of the closest qualifying comparator and the number of eligible comparators, and the position and phase audit of comparisons with ordinary introns.

**Supplementary Table S3.** RNA-seq datasets, junction support, and ordinary intron baseline. Counts are alignment records and can include the same molecule more than once. The last column gives the percentage supported among a fixed random sample of 2,000 ordinary annotated introns per genome. SRR485952 is an EST library.

| Genome | Accession | RNA-seq runs | Library | Primary assayed | Primary supported | Additional assayed | Additional supported | Ordinary introns supported (%) |
|---|---|---|---|---|---|---|---|---|
| *Eimeria tenella* | GCF_000499545.2 | SRR7153211, SRR7153212 | paired RNA-seq | 16 | 5 | 932 | 789 | 77.3 |
| *Eimeria necatrix* | GCF_000499385.1 | SRR14557240 | paired RNA-seq | 28 | 10 | 1,245 | 841 | 66.9 |
| *Eimeria maxima* | GCF_000499605.1 | SRR26671787, SRR26671792 | paired RNA-seq | 14 | 4 | 972 | 767 | 64.6 |
| *Eimeria acervulina* | GCF_000499425.1 | SRR14820228 | paired RNA-seq | 27 | 3 | 1,453 | 999 | 60.2 |
| *Baffinella frigidus* | GCA_026770585.1 | SRR485952 | paired EST | 485 | 439 | 4,727 | 4,134 | 70.3 |
| *Emiliania huxleyi* | GCF_000372725.1 | ERR15561806, ERR15561839, ERR15561871, ERR15561925, SRR37056093, SRR37056094, SRR37056100, SRR37056178 | paired RNA-seq; single RNA-seq | 9 | 7 | 84 | 46 | 31.1 |
| *Symbiodinium* sp. CCMP2456 | GCA_905221635.1 | ERR3711290 | paired RNA-seq | 40 | 30 | 777 | 677 | 90.1 |
| *Symbiodinium* sp. CCMP2592 | GCA_905221615.1 | ERR4501159 | paired RNA-seq | 13 | 11 | 459 | 396 | 81.5 |
| *Symbiodinium* sp. KB8 | GCA_905221625.1 | ERR3711267 | paired RNA-seq | 15 | 12 | 655 | 489 | 75.4 |
| *Symbiodinium natans* | GCA_905221605.1 | SRR32862099 | paired RNA-seq | 156 | 112 | 4,362 | 3,454 | 79.6 |
| *Mesorhabditis spiculigera* | GCA_958295435.1 | ERR11634707, ERR11634708 | paired RNA-seq | 83 | 39 | 1,349 | 541 | 50.3 |
| Total |  |  |  | 886 | 672 | 17,015 | 13,133 | 68.0 |

## Supplementary Figures

**Supplementary Figure S1.**
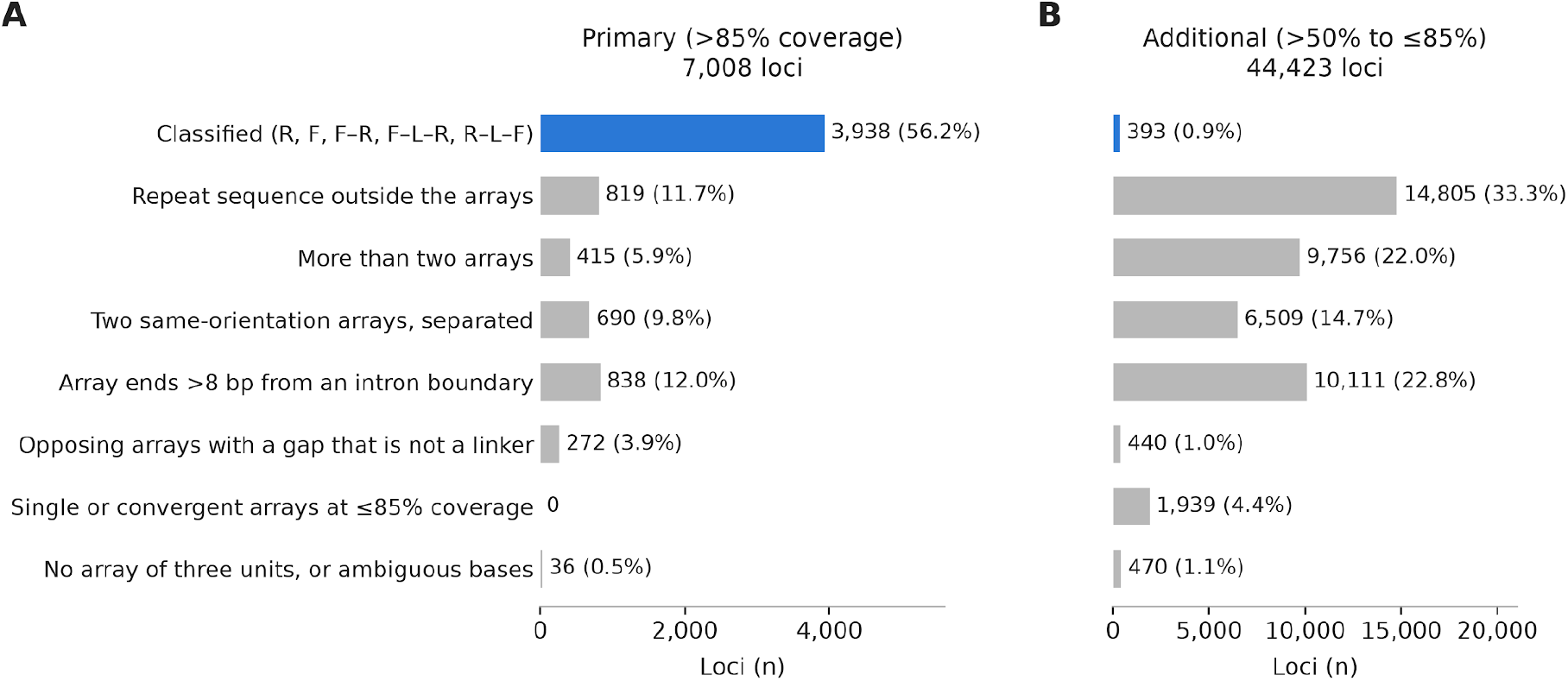
Architecture classification of telotrons. Loci in the primary (A, >85% repeat coverage) and additional (B, >50% to ≤85%) tiers, grouped by the outcome of the architecture classifier. Classified loci (blue) carry one of the five observed architectures. Unclassified loci are grouped by the first rule they fail. The rule each locus failed is in supplementary table S2.

**Supplementary Figure S2.**
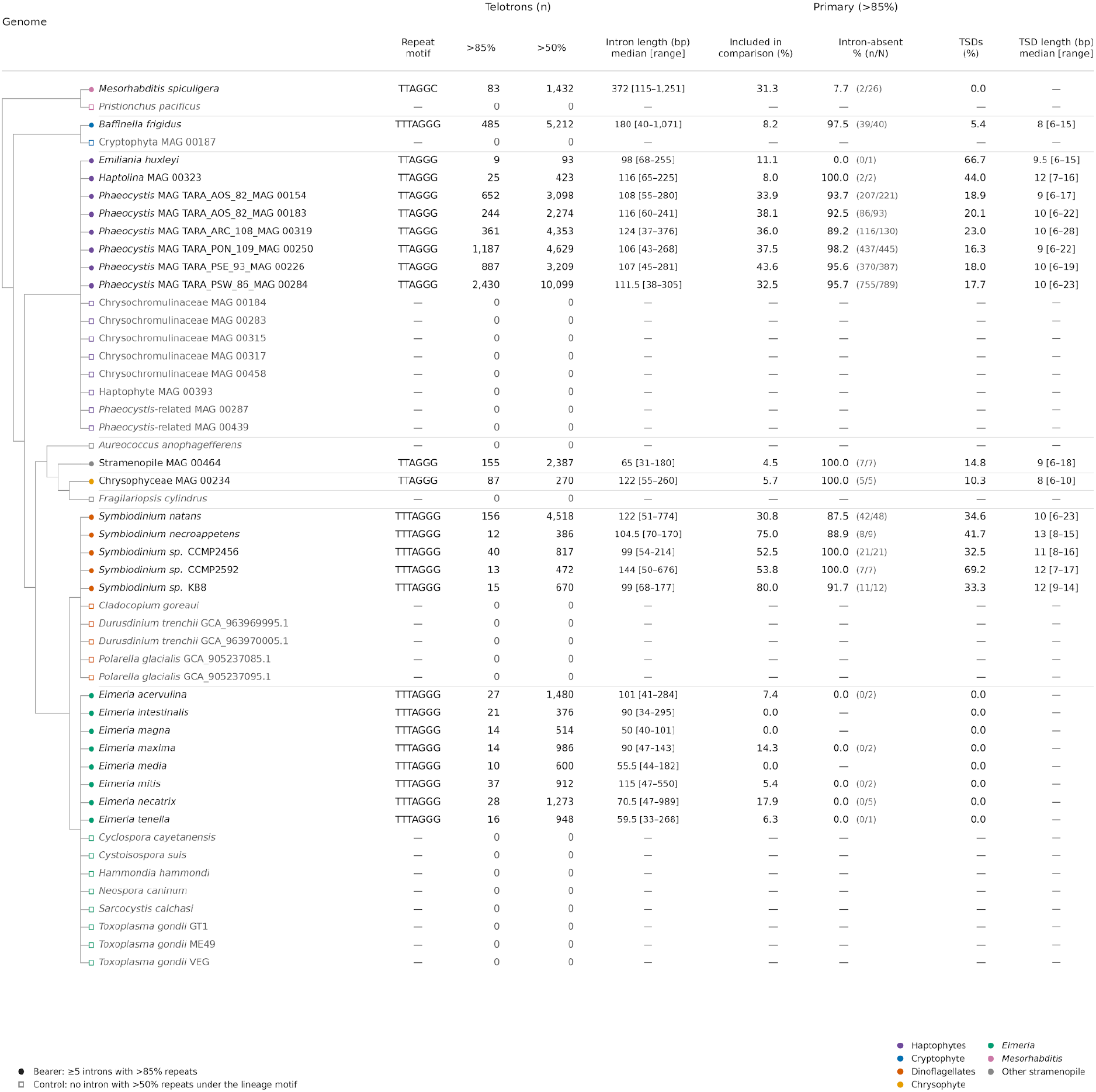
Telotrons and ortholog comparisons in individual genomes. Expanded version of Fig. 1 with the same 25 bearers and 25 controls. The >50% counts include primary loci. Intron and TSD lengths are median [minimum–maximum] in bp. “Included in comparison” is the percentage of primary loci with a supported closest comparison showing an absent intron or an ordinary intron at the same position and phase. The column for absent introns gives absence among those comparisons. TSD percentages use all primary telotron loci in each genome.

**Supplementary Figure S3.**
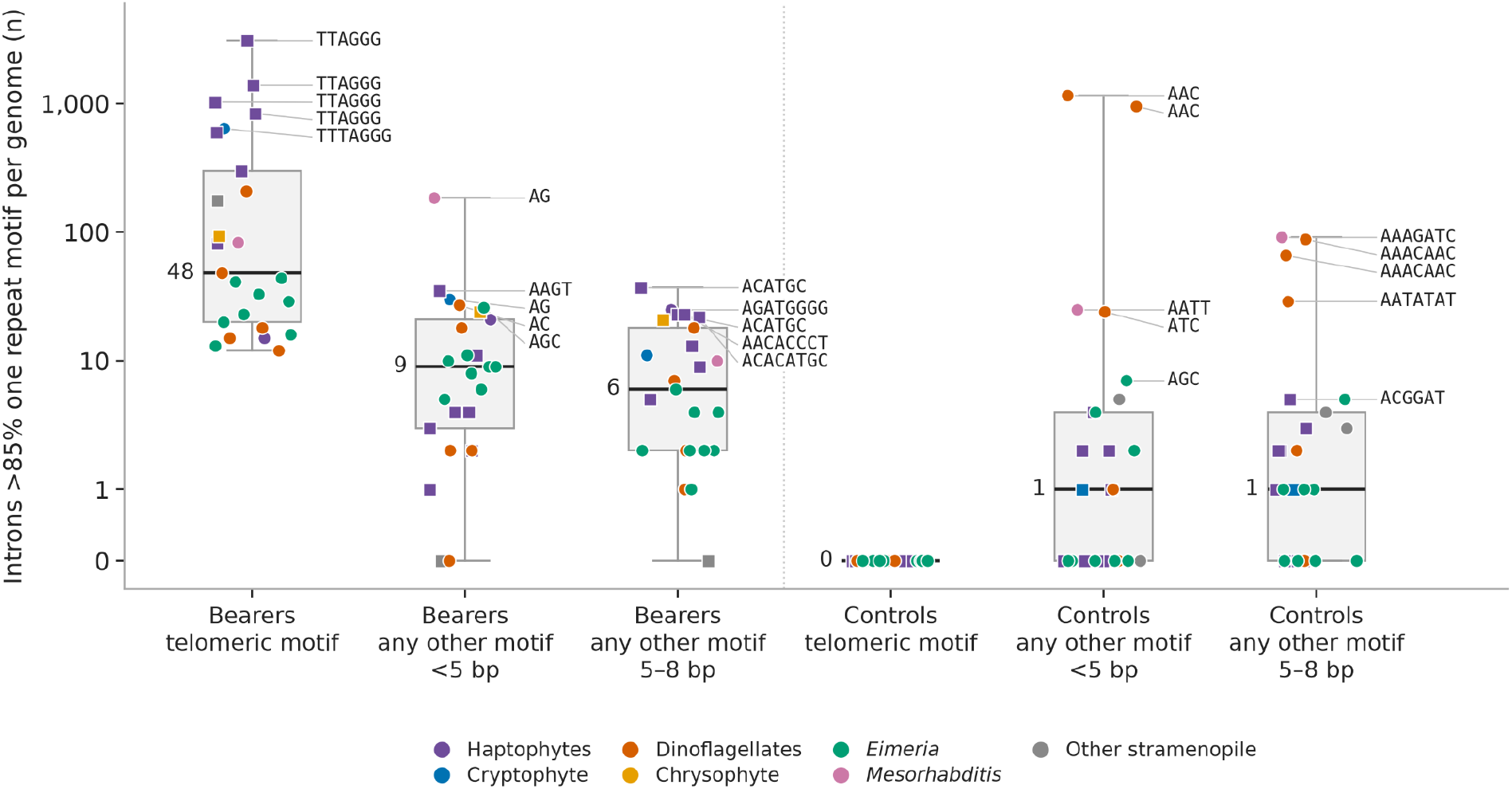
Introns composed of one repeat motif in bearers and controls. Every annotated intron (≥30 bp) of the 25 bearer genomes and their 25 controls was scored against every primitive repeat motif of period 1 to 8 bp. An intron counts once when the motif covering most of it spans more than 85% of its length. Each point is one genome (circles isolates, squares MAGs, colors lineages). Boxes show the median and interquartile range and whiskers show the extremes. The y-axis is logarithmic above 1 and linear below. Telomeric motif: the lineage telomere motif or a motif within one substitution or indel of it. Other motifs are split by period into <5 bp and 5–8 bp. Labels give the motif for the five highest genomes in each group (the lineage motif, or the most frequent other motif). Counts for each bearer genome are in supplementary table S1.

## REFERENCES

Burgt, Ate van der, Edouard Severing, Pierre J. G. M. de Wit, and Jérôme Collemare. 2012. “Birth of New Spliceosomal Introns in Fungi by Multiplication of Introner-like Elements.” Current Biology: CB 22 (13): 1260–1265.

Camacho, Christiam, George Coulouris, Vahram Avagyan, et al. 2009. “BLAST+: Architecture and Applications.” BMC Bioinformatics 10 (December): 421.

Carmel, Liran, Yuri I. Wolf, Igor B. Rogozin, and Eugene V. Koonin. 2007. “Three Distinct Modes of Intron Dynamics in the Evolution of Eukaryotes.” Genome Research 17 (7): 1034–1044.

Choi, Seok-Wan, Louis Graf, Ji Won Choi, et al. 2024. “Ordovician Origin and Subsequent Diversification of the Brown Algae.” Current Biology : CB 34 (4): 740–754.e4.

Collemare, Jérôme, Henriek G. Beenen, Pedro W. Crous, Pierre J. G. M. de Wit, and Ate van der Burgt. 2015. “Novel Introner-Like Elements in Fungi Are Involved in Parallel Gains of Spliceosomal Introns.” PloS One 10 (6): e0129302.

Csuros, Miklos, Igor B. Rogozin, and Eugene V. Koonin. 2011. “A Detailed History of Intron-Rich Eukaryotic Ancestors Inferred from a Global Survey of 100 Complete Genomes.” PLoS Computational Biology 7 (9): e1002150.

Delmont, Tom O., Morgan Gaia, Damien D. Hinsinger, et al. 2022. “Functional Repertoire Convergence of Distantly Related Eukaryotic Plankton Lineages Abundant in the Sunlit Ocean.” Cell Genomics 2 (5): 100123.

Dieterich, Christoph, Sandra W. Clifton, Lisa N. Schuster, et al. 2008. “The Pristionchus Pacificus Genome Provides a Unique Perspective on Nematode Lifestyle and Parasitism.” Nature Genetics 40 (10): 1193–1198.

Gozashti, Landen, Anne Nakamoto, Shelbi Russell, and Russell Corbett-Detig. 2025. “Horizontal Transmission of Functionally Diverse Transposons Is a Major Source of New Introns.” Proceedings of the National Academy of Sciences of the United States of America 122 (21): e2414761122.

Gozashti, Landen, Scott W. Roy, Bryan Thornlow, Alexander Kramer, Manuel Ares Jr, and Russell Corbett-Detig. 2022. “Transposable Elements Drive Intron Gain in Diverse Eukaryotes.” Proceedings of the National Academy of Sciences of the United States of America 119 (48): e2209766119.

Hong, Peihang, Toshihiro Tokiwa, Sijia Yu, and Shyun Chou. 2026. “Polyphyly and Deep Evolutionary History of Piscine Coccidia: Mitochondrial Genomics of a New Goussia Species From Pacific Saury (Cololabis Saira).” The Journal of Eukaryotic Microbiology 73 (5): e70114.

Huff, Jason T., Daniel Zilberman, and Scott W. Roy. 2016. “Mechanism for DNA Transposons to Generate Introns on Genomic Scales.” Nature 538 (7626): 533–536.

Karsenti, Eric, Silvia G. Acinas, Peer Bork, et al. 2011. “A Holistic Approach to Marine Eco-Systems Biology.” PLoS Biology 9 (10): e1001177.

Katoh, Kazutaka, and Daron M. Standley. 2013. “MAFFT Multiple Sequence Alignment Software Version 7: Improvements in Performance and Usability.” Molecular Biology and Evolution 30 (4): 772–780.

Kinzig, Charles G., George Zakusilo, Kaori K. Takai, Logan R. Myler, and Titia de Lange. 2024. “ATR Blocks Telomerase from Converting DNA Breaks into Telomeres.” Science (New York, N.Y.) 383 (6684): 763–770.

Klopfstein, Seraina, and Fredrik Ronquist. 2013. “Convergent Intron Gains in Hymenopteran Elongation Factor-1α.” Molecular Phylogenetics and Evolution 67 (1): 266–276.

Koonin, Eugene V. 2006. “The Origin of Introns and Their Role in Eukaryogenesis: A Compromise Solution to the Introns-Early versus Introns-Late Debate?” Biology Direct 1 (1): 22.

Kumar, Sudhir, Michael Suleski, Jack M. Craig, et al. 2022. “TimeTree 5: An Expanded Resource for Species Divergence Times.” Molecular Biology and Evolution 39 (8). 10.1093/molbev/msac174.

Li, Heng. 2018. “Minimap2: Pairwise Alignment for Nucleotide Sequences.” Bioinformatics 34 (18): 3094–3100.

Li, Heng. 2023. “Protein-to-Genome Alignment with Miniprot.” Bioinformatics (Oxford, England) 39 (1): btad014.

Liu, Hui, Stéphane Aris-Brosou, Ian Probert, and Colomban de Vargas. 2010. “A Time Line of the Environmental Genetics of the Haptophytes.” Molecular Biology and Evolution 27 (1): 161–176.

Li, Wenli, Robert Kuzoff, Chen Khuan Wong, Abraham Tucker, and Michael Lynch. 2014. “Characterization of Newly Gained Introns in Daphnia Populations.” Genome Biology and Evolution 6 (9): 2218–2234.

Li, Wenli, Abraham E. Tucker, Way Sung, W. Kelley Thomas, and Michael Lynch. 2009. “Extensive, Recent Intron Gains in Daphnia Populations.” Science 326 (5957): 1260–1262.

Montresor, Marina, Gabriele Procaccini, and Diane K. Stoecker. 1999. “Polarella Glacialis, Gen. Nov., Sp. Nov. (Dinophyceae): Suessiaceae Are Still Alive!” Journal of Phycology 35 (1): 186–197.

Muyas, Francesc, Manuel José Gómez Rodriguez, Rita Cascão, et al. 2024. “The ALT Pathway Generates Telomere Fusions That Can Be Detected in the Blood of Cancer Patients.” Nature Communications 15 (1): 82.

Nergadze, Solomon G., Mariano Rocchi, Claus M. Azzalin, Chiara Mondello, and Elena Giulotto. 2004. “Insertion of Telomeric Repeats at Intrachromosomal Break Sites during Primate Evolution.” Genome Research 14 (9): 1704–1710.

Nergadze, Solomon G., Marco Andrea Santagostino, Alberto Salzano, Chiara Mondello, and Elena Giulotto. 2007. “Contribution of Telomerase RNA Retrotranscription to DNA Double-Strand Break Repair during Mammalian Genome Evolution.” Genome Biology 8 (12): R260.

Pietluch, Filip, Paweł Mackiewicz Kacper Ludwig, and Przemysław Gagat. 2024. “A New Model and Dating for the Evolution of Complex Plastids of Red Alga Origin.” Genome Biology and Evolution 16 (9). 10.1093/gbe/evae192.

Roy, Scott William. 2016. “How Common Is Parallel Intron Gain? Rapid Evolution Versus Independent Creation in Recently Created Introns in Daphnia.” Molecular Biology and Evolution 33 (8): 1902–1906.

Santagostino, Marco, Francesca M. Piras, Eleonora Cappelletti, et al. 2020. “Insertion of Telomeric Repeats in the Human and Horse Genomes: An Evolutionary Perspective.” International Journal of Molecular Sciences 21 (8): E2838.

Schrumpfová, Petra Procházková, and Jiří Fajkus. 2020. “Composition and Function of Telomerase-A Polymerase Associated with the Origin of Eukaryotes.” Biomolecules 10 (10). 10.3390/biom10101425.

Sola, Lorenzo, Solomon G. Nergadze, Eleonora Cappelletti, Francesca M. Piras, Elena Giulotto, and Marco Santagostino. 2021. “Telomeric-like Repeats Flanked by Sequences Retrotranscribed from the Telomerase RNA Inserted at DNA Double-Strand Break Sites during Vertebrate Genome Evolution.” International Journal of Molecular Sciences 22 (20): 11048.

Sunagawa, Shinichi, Silvia G. Acinas, Peer Bork, et al. 2020. “Tara Oceans: Towards Global Ocean Ecosystems Biology.” Nature Reviews. Microbiology 18 (8): 428–445.

Torii, Kota, Chihiro Ota, Ayumi Nagashima, Masaki Kajikawa, and Akira Kato. 2025. “Intron Turnover of slc26a1 and slc26a2 and Convergence of Intron Insertion Sites.” Scientific Reports 15 (1): 30007.

Worden, A. Z., J. H. Lee, T. Mock, et al. 2009. “Green Evolution and Dynamic Adaptations Revealed by Genomes of the Marine Picoeukaryotes Micromonas.” Science 324 (5924): 268–272.

Yenerall, Paul, and Leming Zhou. 2012. “Identifying the Mechanisms of Intron Gain: Progress and Trends.” Biology Direct 7 (1): 29.

